# Climatic stability and resource availability explains dung beetles (Scarabaeinae) richness patternon the Americas

**DOI:** 10.1101/402537

**Authors:** Anderson Matos Medina

**Affiliations:** Programa de Pós□Graduação em Ecologia e Evolução, Instituto de Ciências Biológicas, Universidade Federal de Goiás, Goiânia, Goiás, Brazil

**Keywords:** climatic instability, resource diversity, dung beetle diversity, mammal richness, Scarabaeidae

## Abstract

Climatic conditions are the main driver of species richness. Specifically, the increase in climatic instability may reduce species richness directly and indirectly by reducing resources available. This hypothesis is evaluated here using a producer-consumer interaction to explain dung beetle richness on a continental scale (America) using mammal richness as resource proxy and temperature and precipitation seasonality as a proxy for climatic instability. A spatial path analysis was built in order to evaluate this hypothesis while controlling for spatial autocorrelation and differences in the sampling effort and abundance of each study (n=115) gathered from the literature. Dung beetle richness was directly explained by temperature seasonality, precipitation seasonality, and mammal richness, whereas only precipitation seasonality had an effect modulated by mammal richness. This result reinforces the notion that species richness can be explained by climatic conditions, but also reveals the importance of biotic interactions in order to understand the mechanisms behind such patterns.

## Introduction

Latitudinal diversity gradients are a widespread pattern in ecology (Gaston 2000; Willig et al. 2003; Hillebrand 2004). To understand latitudinal gradient one must go beyond the pattern and therefore evaluate causal hypotheses, which encompass three types of explanations: ecological, historical and evolutionary (Gaston 2000; Hawkins and Diniz-Filho 2004; Mittelbach et al. 2007). Examples of hypotheses proposed for this geographic pattern are productivity, environmental heterogeneity, area, historical factors and biotic interactions (Willig et al. 2003; Field et al. 2009). Among the above mentioned, climatic hypotheses are between the most used explanations behind the pattern of latitudinal diversity gradients (Field et al. 2009). However, climatic stability can have different weights in determining the latitudinal gradient. One rationale is climatic stability is more important to determine the latitudinal gradient in temperate regions (e.g.: temperate species have higher climatic tolerance than their tropical counterparts (Stevens 1989)), whereas biotic interactions play a major role in tropical regions (Dobzhansky 1950). Albeit this is not a new idea, our understanding of how species richness changes along latitudinal gradients can be improved by weighting the relative contributions of climatic stability and biotic interactions.

The aggregate effects of climatic stability and biotic interactions can be noticed, for example, when variations on temperature and rainfall can reduce the time window when resources are available for species which may generate resource bottlenecks that can limit the number of species coexisting (Williams and Middleton 2008). In fact, the climatic stability hypothesis predicts that sites with higher stability possesses higher richness than sites with lesser stability (Pianka 1966), and one of the explanations for such phenomena invokes the role of biotic interactions because climatic stability would allow species to be more specialized on stable environments (Moles and Ollerton 2016; but see Schleuning et al. 2012). Direct effects of resource availability on species can also play a role in explaining the causes of the latitudinal richness pattern. For example, the number of species of woody plants has been used to explain the richness of arboreal or frugivorous mammals, and figs richness has been demonstrated to determine frugivorous birds richness (Andrews and O’Brien 2000, Kissling et al. 2007; but see Hawkins and Pausas 2004). The rationale behind this hypothesis is that an increase in species richness at basal trophic levels should increase the number of species at higher trophic levels.

Here, I focus on how dung beetle diversity pattern can be shaped by biotic interactions between fecal detritus producers (mammals) and consumers (dung beetles) (Nichols et al. 2016) along with climatic stability. Dung beetles (Scarabaeinae) is a rich group of detritivores beetles that probably diversified after the transition from saprophagy to coprophagy (Halffter 1991). In this study, two hypotheses will be evaluated. First, if resource availability is important for richness patterns of dung beetles then it is expected that dung beetles diversity should be driven by mammals’ diversity since mammals dungs are the primordial source of food resource for dung beetles (Halffter and Matthews 1966; Nichols et al. 2009). Examples of how dung beetle diversity is related to mammal diversity are that hunting-related reductions on mammals richness lead to a reduction in dung beetles richness (Nichols et al. 2009; Culot et al. 2013) and that changes on mammals species composition may explain changes on dung beetle composition (Bogoni et al. 2016). Second, if variations on climatic stability should affect dung beetles then I expect to find that an increase in climatic stability should have positive effects on dung beetle richness. In general, many insects display seasonality patterns of diversity (Wolda 1988), and dung beetle communities, in particular, have increased richness with increased temperature and reduction of rainfall seasonality (Andresen 2005; Hernández and Vaz-de-Mello 2009; Liberal et al. 2011). Climatic stability should also allow dung beetles species to be more specialized in feeding resources, therefore an increase in niche partitioning between co-occurring dung beetle species caused by trophic specialization could increase dung beetle species richness (Larsen et al. 2006), especially in tropical areas where higher mammal richness are also expected (Davies and Buckley 2011; Safi et al. 2011). Even though most dung beetles are generalists and only a few species can be sorted in guilds that use different dung types (Filgueiras et al. 2009; Bogoni and Hernandez 2014).

In this study, I test the hypothesis that dung beetle latitudinal richness is explained by 1) increase in feeding resources availability because of biotic interactions with mammals that produce dung beetle feeding resources, and 2) an increase on climatic stability and 3) both resource availability and climatic stability. To evaluate these hypotheses, I used a spatialized path analysis which allows dissociating the direct and indirect effects of the climatic stability. First, I predict that dung beetle richness should increase due increases in climatic stability, in other words, decreases in precipitation seasonality and temperature seasonality. Second, increases on mammals’ richness, here used as a proxy for resource availability, should result in increases of dung beetle richness. Since mammals richness can also respond to precipitation seasonality and temperature seasonality, therefore climatic stability indirect effects can change dung beetle richness through mediated effects in mammals richness (See Figure 1 for more details on proposed path analysis).

**Figure 1.**
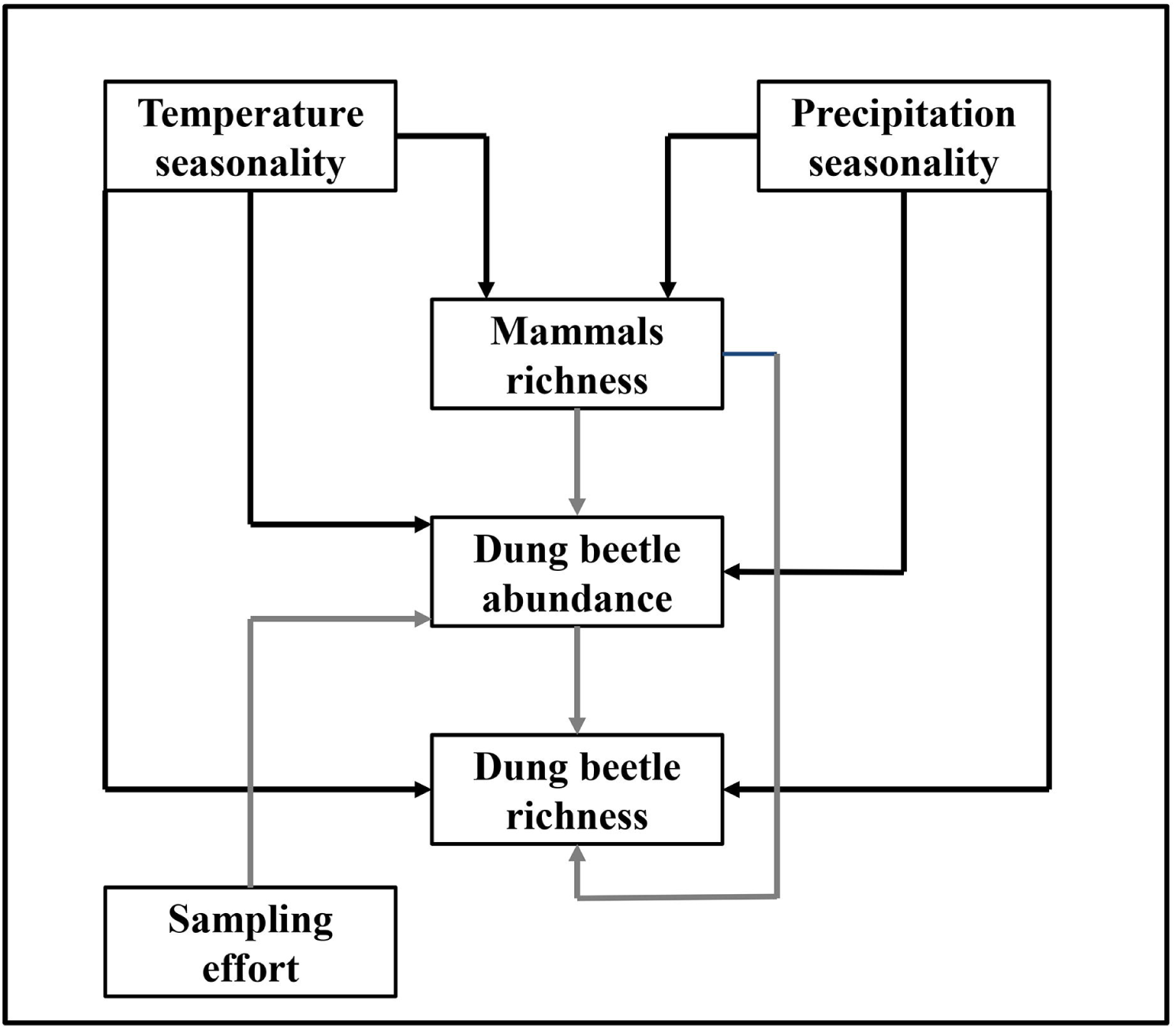
Path analysis representing the theoretical relationships between predictors of dung beetle richness. Black arrows are negative effects while gray arrows are positive relationships.

## Materials and Methods

### Dung beetle database

I performed a search on the Scopus database for dung beetles inventories using the following keywords: “*dung beetle”* or “*Scarabaeinae”*. I only included the period between 1980 and 2016. This search retrieved a total of 1443 articles and after applying the following criteria, the database comprised 115 studies and 213 sites (Figure 2): 1) sampled dung beetles should belong to the subfamily Scarabaeinae (in other words, studies with only Aphodinae, Geotrupinae and Troginae were excluded); 2) the study employed a standardized sampling protocol and clearly stated the number of pitfalls, sites and temporal replication used while sampling the dung beetles. Sampling protocol information was used to the measure sampling effort using in each study. The sampling effort was the total number of pitfalls measured as pitfalls multiplied per area multiplied per temporal replications. Differences on use of different baits (types of dung, decomposing meat or rotten fruits) were ignored 3) provided a list of sampled species and their respective abundances 4) sampled in the American continent and presented geographic coordinates or specified the municipality where the sampled occurred. If one study sampled more than one site and provided a geographic coordinate for each site thus each site was included on the database. However, species lists were combined in cases where there was only one geographic coordinate for multiples sites. Studies included in the analysis are available on the Supplementary File 1.

**Figure 2.**
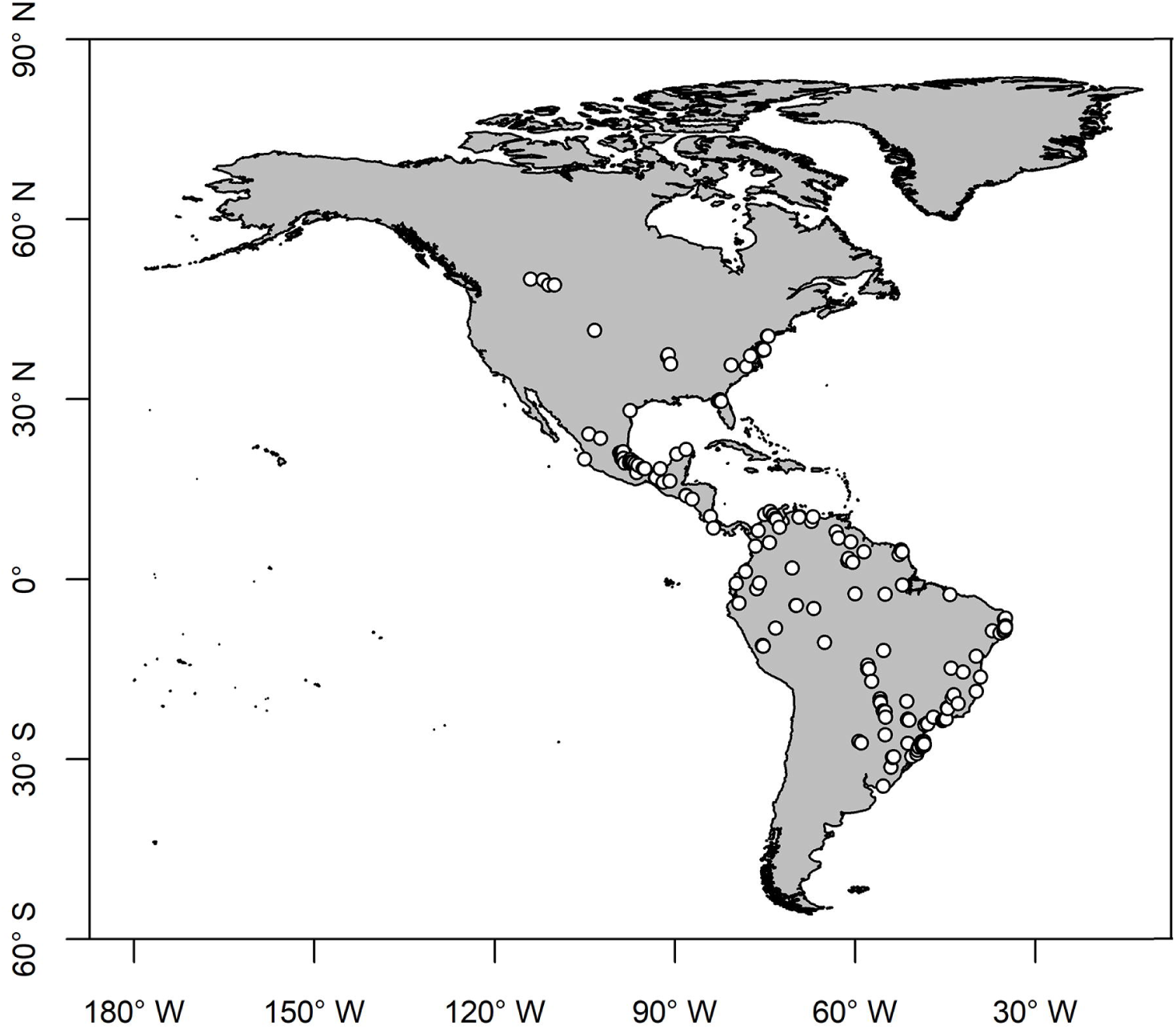
Localities of the 115 dung beetle inventories performed on the America continent and used on the analysis.

### Explanatory variables

I used mammal richness as a proxy for resource availability because of biotic interactions between dung beetles and mammals (Nichols et al. 2009, 2013). In order to measure mammal richness, I downloaded the terrestrial mammals’ red list range shapefile (IUCN 2016) to estimate the number of mammals richness in the locations where dung beetles were sampled. For each dung beetle locality, I counted the number of mammal species’ range shapefiles that overlap their coordinates using package *maptools* (Bivand and Lewin-Koh 2017).

Climatic stability was measured as precipitation seasonality and temperature seasonality obtained from the worldclim with 5 arc-minutes resolution (approximately 10 km at equator line) (Hijmans et al. 2005). I opted for a coarser resolution because many localities had their geographic coordinates manually assigned. These variables are important drivers of mammals distribution and are commonly used on studies with species distribution of mammals (e.g.: Moura et al. 2016, Ribeiro et al. 2016). Values of precipitation seasonality and temperature seasonality were extracted for each locality using package raster (Hijmans 2016).

Both sampling effort and dung beetle abundance were used to control the effects of different sampling protocols and sampling efforts on dung beetle richness because measures of species richness disregarding differences in sampling effort may lead to biases estimations (Gotelli and Colwell 2001). Additionally, dung beetle abundance may change positively to increased climatic stability and resource stability because of other hypotheses (e.g.: more individuals hypothesis), therefore these confounding effects are minimized by incorporating this path in the structural model equations. Furthermore, there are cases in which dung beetle abundance may increase with a decrease in mammals’ richness (Culot et al. 2013). However, this is an effect of selective defaunation of large mammals and if this effect is noticeable it should be accounted in the path analysis.

### Data Analysis

A path analysis was built using the piecewise structural equation modeling approach that allows the incorporation of different models by building each model separately (Shipley 2009; Lefcheck 2016). Additionally, path analysis helps to disentangle direct and indirect effects of climatic and resources on the richness patterns (Kissling et al. 2007; Moura et al. 2016). First, I built three generalized least squares models (GLS) in order to account spatial autocorrelation using the package *nlme* (Pinheiro et al. 2014). Spatial correlation structure present on the three models was reduced by selecting the best among four spatial correlations: Spheric, Exponential, Gaussian and Rational quadratic. The first model was built using mammal richness as the response variable, precipitation seasonality and temperature seasonality as explanatory variables, and using a rational quadratic spatial correlation structure. The second model was built using dung beetle abundance as the response variable, precipitation seasonality, temperature seasonality, mammal richness and sampling effort as explanatory variables, and using an exponential spatial correlation structure. The third model was built using dung beetle richness as the response variable and precipitation seasonality, temperature seasonality, mammal richness and dung beetle abundance as explanatory variables. These three models were assembled in a path analysis using the package *piecewiseSEM* (Lefcheck 2016). The significance of the piecewise SEM was evaluated using Fisher’s C statistic in which a p-value above 0.05 is an indication that the model fits well to the data.

In all models, dung beetle abundance and dung beetle richness were log10 transformed in order to achieve residuals with a normal distribution. All analyses were carried out on the R environment (R Core Team 2017). More details in model selection and models assumptions are found in the Supplementary file 2.

## Results

Dung beetle richness ranged from one to 101 species (mean ± SD = 21.5 ± 17.2), whereas mammals richness ranged from three to 193 species (113.9 ± 39.5). Dung beetle abundance ranged from seven to 93,274 individuals (5,226.8 ± 11,606.66).

I found that the proposed path analysis had a good fit to the observed data (Fisher’s C = 5.05; d.f. =4; P=0.282; Figure 3). Mammal richness was negatively affected both by temperature seasonality and precipitation seasonality (R^2^=71.31%) whereas dung beetle abundance variation was explained only by sampling effort (R^2^=9.03%).

**Figure 3.**
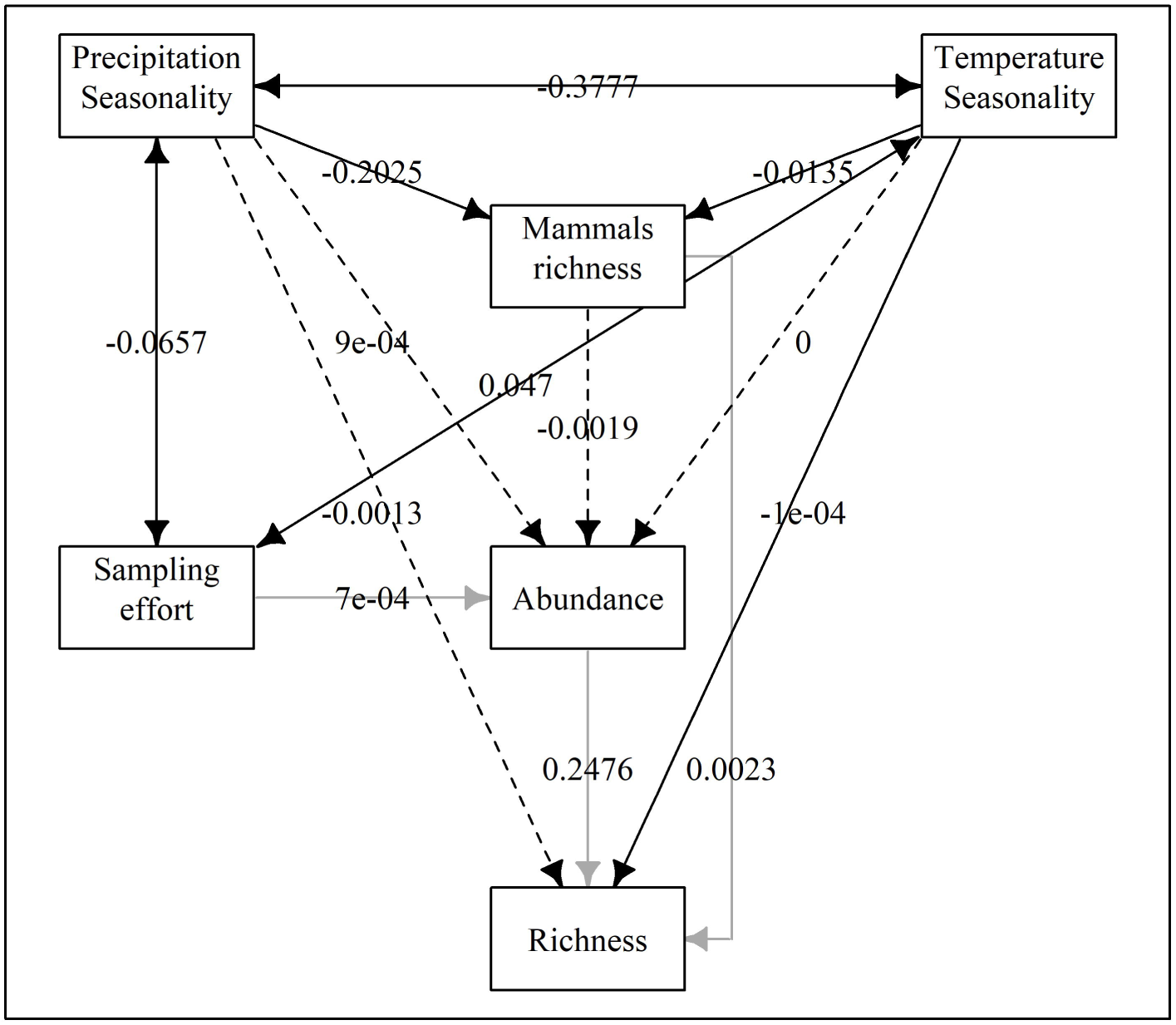
Observed path analysis used to predict dung beetle richness. Dashed arrows are non-significant relationships between variables (p>0.05) while continuous arrows are significant relationships (p<0.05). Black arrows are negative effects while dark gray arrows are positive effects. Double arrow represents correlations between exogenous variables. Note that both dung beetle richness and dung beetle abundance was log transformed.

In the model, 56.32% of the variation on dung beetle richness was explained. An increase on four units of dung beetle abundance resulted on an increase of one unit on dung beetle richness (b = 0.2472), and an increase of mammal richness resulted on an increase of dung beetle richness (b_log10_=0.0023; b_antilog_= 1.01). As expected, negative effects on dung beetle richness included a direct and an indirect effect, modulated through mammal richness, of temperature seasonality (b_direct_=-0.0001and b_indirect_= - 0.00003), but only indirect effects of precipitation seasonality dung beetle richness (b_indirect_= −0.0005).

## Discussion

Here I have corroborated my hypothesis that dung beetle richness was higher in areas with higher climatic stability and higher resource availability. Despite this, dung beetle abundance was poorly explained on the model and that could be a consequence of lack of standardization between sampling protocols of studies (Larsen and Forsyth 2005; da Silva and Hernández 2015). As far as I have know, this is among the firsts empirical assessments of dung beetle richness pattern (see: Frank et al. 2018), since only anecdotal evidence and local scale assessments were previously made (Halffter and Matthews 1966; Hernández and Vaz-de-Mello 2009; Liberal et al. 2011).

Two non-exclusive explanations help to understand why there are more dung beetle species on areas with more resources available (i.e. higher mammals’ richness). First, an increase on resource availability should allow dung beetles to specialize on certain types of dung (e.g.: Larsen et al. 2006, Jacobs et al. 2008) and reduce interspecific competition between dung beetles. However, a recent study has shown an opposite trend – dung beetles have low specificity on resource use even on communities with high richness (Frank et al. 2018). Second, an increase in mammal richness could result in an increase of different traits and lineages of mammals (Safi et al. 2011). That, in turn, can increase the number of mammals with different activity times (e.g.: diurnal or nocturnal) and with different feeding and digestive systems (e.g.: herbivores or carnivores). Indeed, dung beetle community structure changes between night and day (Lopes et al. 2011) or depending on dung type used as bait (Filgueiras et al. 2009).

Climatic instability affects directly the time of activity when dung beetles can be active due to physiological constraints, mostly of small dung beetles (<2g) that are thermoconformers (Verdú et al. 2006). Considering that areas with more climatic instability should have a higher variation on temperature during the day and during the year, this could in turn limit the number of dung beetles species active. Additionally, climatic instability should also increase the number of generalist dung beetle species in order to deal with a decrease on resource availability, climatic instability also reduces the number of mammals’ species, which reduces dung beetle species due to interspecific competition (Finke and Snyder 2008). Surprisingly, dung beetle richness was not directly affected by precipitation seasonality. This once again points towards climatic instability having an indirect effect through resource availability.

Here I had shown the importance of incorporating biotic interactions, here as trophic relationships between dung beetles and mammals, in order to decouple direct and indirect effects of climatic on species richness patterns, particularly because patterns of trophic groups can arise from random effects (Gaston 2000). For example, a study revealed that congruent pattern of richness found with ants and trees are mostly explained by similar responses to environmental gradients (Vasconcelos et al. 2019). Furthermore, I show analytical evidence that dung beetle richness is largely affected by mammals’ richness and that mammal’s richness can mediate climatic instability effects on dung beetles. My finding increases the understanding of the intrinsic relationship between detritivore (dung beetle) and producers (mammals) which have an important role in ecosystem services (e.g.: nutrient cycling and seed dispersal) (Nichols et al. 2009) and since dung beetle are widely dependent of mammals conservation actions must take in consideration the need to preserve the actors involved in the detritus food web in order to conserve these ecosystem services.

## Acknowledgments

I received a scholarship from Coordenação de Aperfeiçoamento de Pessoal de Nível Superior (CAPES). I am also grateful for the effort of all authors that sampled dung beetle communities used as source data for this study.

