## Supplementary File 1 for "Climatic stability and resource availability explains dung beetles (Scarabaeinae) richness patternon the Americas"

Title: Following the dung or the climate? Species richness pattern of dung beetles (Scarabaeinae) on the Americas

Journal of Insect Conservation

Author: Anderson Matos Medina

Supplementary Information 1: Source data and duplicate removal

Table 1. List of studies that had at least one duplicates site and the criteria used to remove those studies. Priority was given to studies that were published more recently, had more sampling effort, had abundance information and did not included flight intercept sampling.

| Used | Removed | Criteria |
| --- | --- | --- |
| Basto-Estrella et al., 2014 | Basto-estrella et al., 2012 | more recent |
| Campos & Hernández, 2015b | Campos & Hernández, 2015a | more effort |
|  | Campos & Hernández, 2013 | more effort |
| Costa et al., 2013 | Iannuzzi et al., 2016 | more effort |
| da Silva & Hernández, 2015 | da Silva & Hernández, 2014 | more effort |
|  | da Silva & Hernández, 2016 | more effort |
| da Silva et al., 2012a | Audino et al., 2011 | pitfall + flight |
| Feer & Boissier, 2015 | Price & Feer, 2012 | more points |
|  | Feer, 2013 | more points |
| Hernández & Vaz-de-Mello, 2009 | Hernández et al., 2011 | Abundance |
| Horgan, 2001 | Horgan, 2002 | more points |
| Korasaki et al., 2013b | Lopes et al., 2011 | more points |
| Medina & Lopes, 2014a | Medina & Lopes, 2014b | abundance |
| Price, 2004 | Price, 2006 | more points |
| Rangel-Acosta et al., 2016a | Hernandez et al., 2012 | more effort |
| Silva et al., 2010 | Costa et al., 2009 | pitfall + flight |
| Verdú et al., 2007 | Barragán et al., 2011 | Abundance |
