## Supplementary File 2 for "Climatic stability and resource availability explains dung beetles (Scarabaeinae) richness patternon the Americas"

Title: Following the dung or the climate? Species richness pattern of dung beetles (Scarabaeinae) on the Americas

Author: Anderson Matos Medina

Supplementary Information 2: Model selection and model assumption

For each response variable (abundance, richness and mammals’ richness), four generalized least squares (GLS) were built with a spatial autocorrelation structure and one GLS without any assumption of spatial autocorrelation structure (*Raw*). Models were compared using AIC (Burnham and Anderson 2002) and the model with the lesser amount of AIC was selected as the best model (Zuur et al. 2009). Each of the best model (abundance, richness and mammals’ richness) was used to in the piecewise structural equation modeling (SEM).

Table S1. Model selection of spatial correlation structure for each response variable.

| Response | Correlation | df | AIC | ΔAIC |
| --- | --- | --- | --- | --- |
| Abundance | Exponential | 8 | 424.43 | 0 |
|  | Ratio | 8 | 426.58 | 2.15 |
|  | Gaussian | 8 | 428.34 | 3.92 |
|  | Spherical | 8 | 454.72 | 30.29 |
|  | Raw | 6 | 518.94 | 94.51 |
| Richness | Exponential | 8 | -33.40 | 0 |
|  | Gaussian | 8 | -23.81 | 9.58 |
|  | Spherical | 8 | -22.70 | 10.70 |
|  | Ratio | 8 | -12.73 | 20.66 |
|  | Raw | 6 | 37.91 | 71.31 |
| Mammals’ richness | Ratio | 6 | 1640.48 | 0 |
|  | Gaussian | 6 | 1662.09 | 21.61 |
|  | Exponential | 6 | 1683.86 | 43.38 |
|  | Spherical | 6 | 1702.98 | 62.50 |
|  | Raw | 4 | 1996.60 | 356.12 |

Multicollinearity of the data was evaluated by measuring variance inflation factor (VIF) separately for the best generalized least square model selected for each regression model (GLS considering a spatial autocorrelation class) using package *car* (Fox and Weisberg 2011). All variables had VIF values below 2.5 (Table S2).

Table S2. Variance inflation factor measured for the best-fitted model.

| Response | Precipitation seasonality | Temperature  seasonality | Mammal  richness | Abundance | Sampling  Effort |
| --- | --- | --- | --- | --- | --- |
| Abundance | 1.13 | 2.33 | 2.16 | - | 1.01 |
| Richness | 1.09 | 1.69 | 1.60 | 1.02 | - |
| Mammal richness | 1.06 | 1.06 | - | - | - |

For each model created, spatial autocorrelation was inspected using Moran’s I correlogram built using distance classes with increasing distance (800m) using package *ncf* (Bjornstad 2016). Small spatial correlation (Moran’s I between -0.2 and 0.2) remained on all models. Furthermore, selected models had moderate levels of autocorrelation on the last classes for abundance (Fig S1) and richness (Fig S2), however, mammal richness selected model had a high level of spatial autocorrelation for the last distance class (Fig S3).

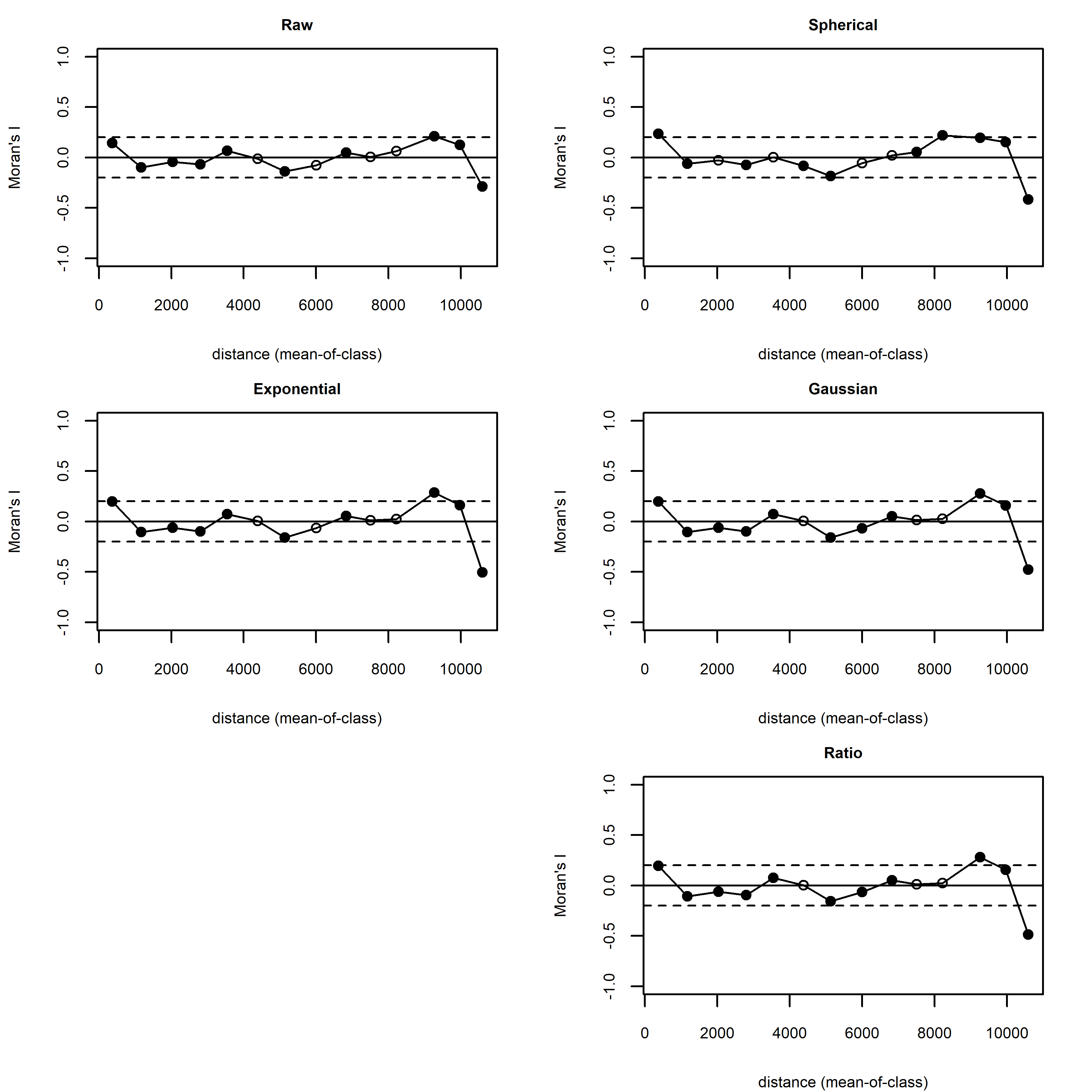

Figure S1. Moran’s I correlograms from the residuals from the five models built for explaining dung beetle abundance. White circles are non-significant spatial correlations while black circles are significant correlations. Dashed lines correspond to the [-0.2, 0.2] interval. Spherical spatial autocorrelation model was selected.

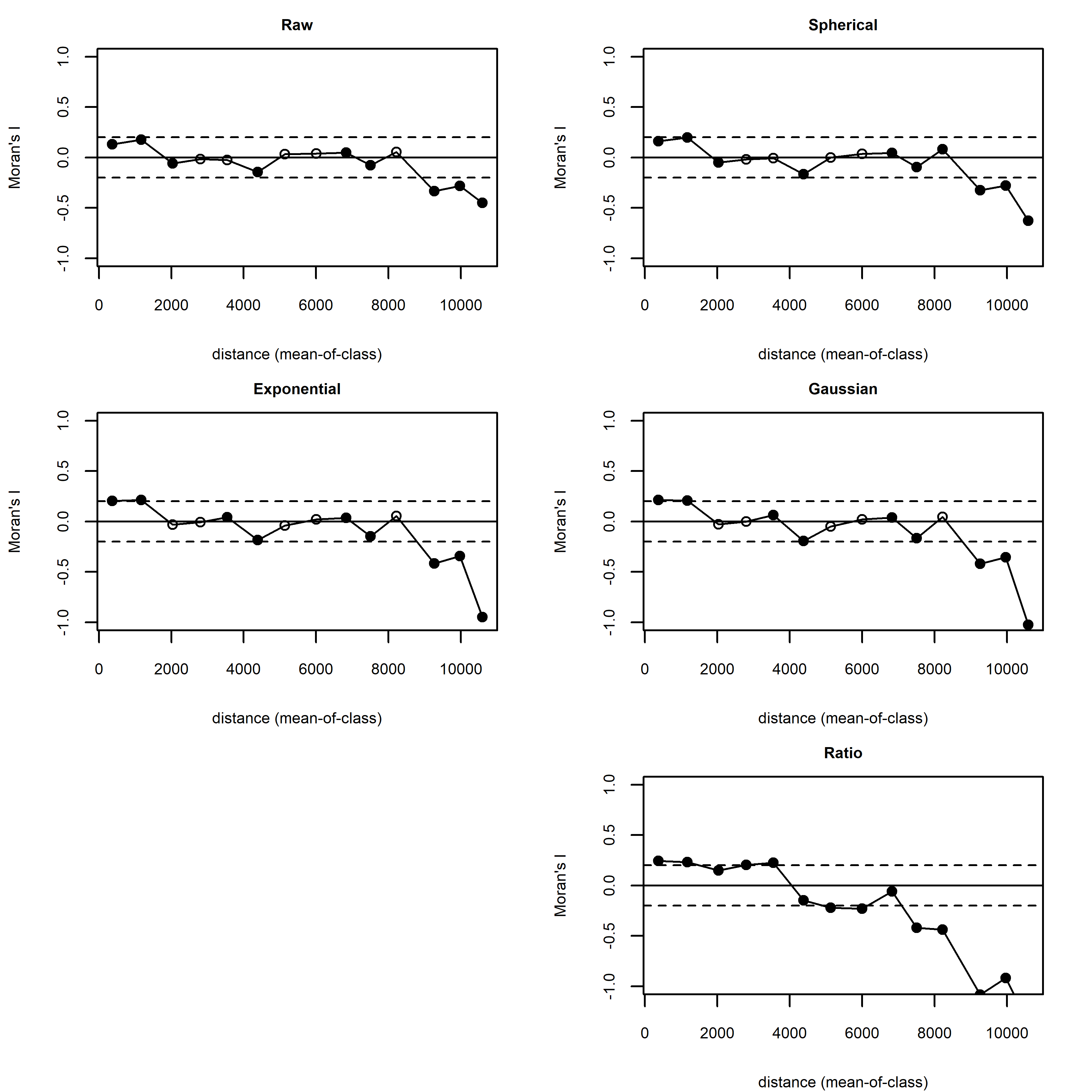

Figure S2. Moran’s I correlograms from the residuals from the five models built for explaining dung beetle richness. White circles are non-significant spatial correlations while black circles are significant correlations. Dashed lines correspond to the [-0.2, 0.2] interval. Spherical spatial autocorrelation model was selected.

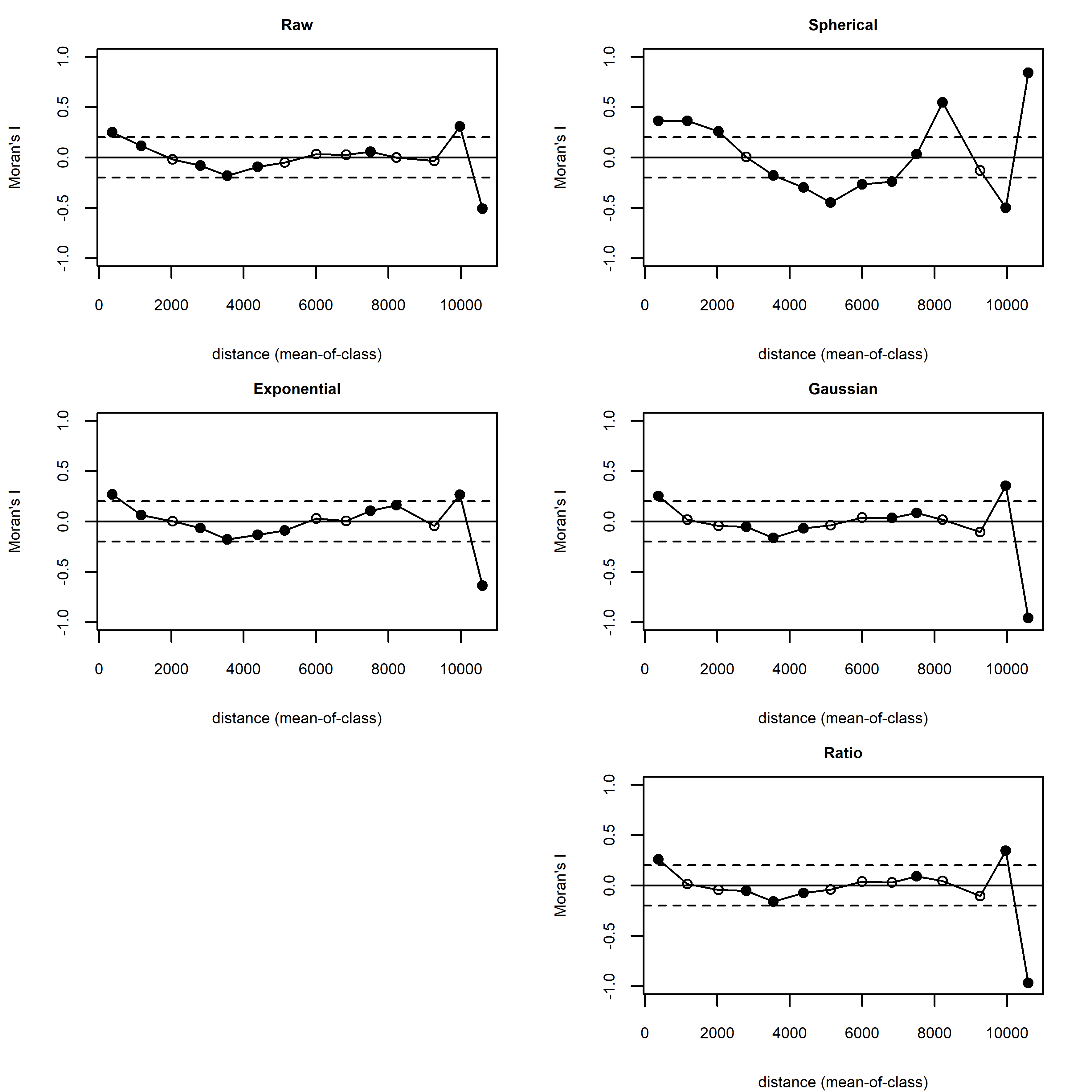

Figure S3. Moran’s I correlograms from the residuals from the five models built for explaining mammals richness. White circles are non-significant spatial correlations while black circles are significant correlations. Dashed lines correspond to the [-0.2, 0.2] interval. Ratio spatial autocorrelation model was selected.
